# Insights into molecular evolution recombination of pandemic SARS-CoV-2 using Saudi Arabian sequences

**DOI:** 10.1101/2020.05.13.093971

**Authors:** Islam Nour, Ibrahim O. Alanazi, Atif Hanif, Alain Kohl, Saleh Eifan

## Abstract

The recently emerged SARS-CoV-2 (*Coronaviridae; Betacoronavirus*) is the underlying cause of COVID-19 disease. Here we assessed SARS-CoV2 from the Kingdom of Saudi Arabia alongside sequences of SARS-CoV, bat SARS-like CoVs and MERS-CoV, the latter currently detected in this region. Phylogenetic analysis, natural selection investigation and genome recombination analysis were performed. Our analysis showed that all Saudi SARS-CoV-2 sequences are of the same origin and closer proximity to bat SARS-like CoVs, followed by SARS-CoVs, however quite distant to MERS-CoV. Moreover, genome recombination analysis revealed two recombination events between SARS-CoV-2 and bat SARS-like CoVs. This was further assessed by S gene recombination analysis. These recombination events may be relevant to the emergence of this novel virus. Moreover, positive selection pressure was detected between SARS-CoV-2, bat SL-CoV isolates and human SARS-CoV isolates. However, the highest positive selection occurred between SARS-CoV-2 isolates and 2 bat-SL-CoV isolates (Bat-SL-RsSHC014 and Bat-SL-CoVZC45). This further indicates that SARS-CoV-2 isolates were adaptively evolved from bat SARS-like isolates, and that a virus with originating from bats triggered this pandemic. This study thuds sheds further light on the origin of this virus.

**AUTHOR SUMMARY:** The emergence and subsequent pandemic of SARS-CoV-2 is a unique challenge to countries all over the world, including Saudi Arabia where cases of the related MERS are still being reported. Saudi SARS-CoV-2 sequences were found to be likely of the same or similar origin. In our analysis, SARS-CoV-2 were more closely related to bat SARS-like CoVs rather than to MERS-CoV (which originated in Saudi Arabia) or SARS-CoV, confirming other phylogenetic efforts on this pathogen. Recombination and positive selection analysis further suggest that bat coronaviruses may be at the origin of SARS-CoV-2 sequences. The data shown here give hints on the origin of this virus and may inform efforts on transmissibility, host adaptation and other biological aspects of this virus.

## INTRODUCTION

A novel human pathogen called severe acute respiratory syndrome coronavirus 2 (SARS-CoV-2; *Coronaviridae; Betacoronavirus*) originated from Hubei, China in December 2019 and has since spread all around the world [1]. The disease was named as COVID-19 and human to human transfer has been established [2]. The disease symptoms depicted in SARS-CoV-2 infections were found similar to the infections caused by SARS coronavirus (SARS-CoV) in 2003 [3], however it would appear that the case case fatality rate is considerably lower [4]. A virus related to SARS-CoV, Middle East Respiratory syndrome coronavirus (MERS-CoV) originated from camels in the Middle East and cases are still reported by the Ministry of Health of the Kingdom of Saudi Arabia [5, 6].

SARS-CoV-2 is different from two zoonotic coronaviruses, SARS-CoV and MERS-CoV that caused human disease earlier in the twenty-first century. Beforehand, the *Coronaviridae* Study Group, an ICTV working group, determined each of these later two viruses prototype as a new species in new subgenera of the genus *Betacoronavirus*, *Sarbecovirus* and *Merbecovirus* [7, 8]. SARS-CoV-2 was assigned recently to the sarbecoviruses, a grouping that contains hundreds of known viruses predominantly isolated from humans and diverse bats [9].

Coronaviruses are positive sense, non-segmented, single stranded, enveloped RNA viruses with genome size of 26 kb to 32 kb identified to cause respiratory diseases in a variety of animals and humans. Human coronaviruses like SARS-CoV, MERS-CoV, and SARS-CoV-2 are pathogens of zoonotic origin [10]. Previous sequence analysis showed a high percentage of similarity among SARS-CoV-2, SARS-CoV and bat corona viruses [11, 12].

Coronaviruses contains mainly four types of structural and several non-structural proteins [10, 13, 14]. The spike protein S is one of the structural protein plays a key role in recognition and attachment of SARS-CoV and SARS-CoV-2 to the host cell angiotensin-converting enzyme 2 (ACE2) receptor [15, 16, 17]. Structurally, S is composed of two functional subunits essential for binding to the host cell receptor (S1 subunit) and virus-host cell fusion (S2 subunit) [18]. The S1 subunit exists within the N-terminal 14–685 amino acids of S, including the N-terminal domain (NTD), receptor binding domain (RBD), and receptor binding motif (RBM). The S2 subunit involves fusion peptide (FP), heptad repeat 1 (HR1), heptad repeat 2 (HR2), transmembrane domain I and cytoplasmic domain (CP). Moreover, SARS-CoV-2 S protein comprises a special S1/S2 furin-detectible site, leading to potentially distinctive infectious properties (12). SARS-CoV-2 genome analysis depicted a similarity index of 79.5% with SARS-CoV and very high resemblance to bat coronaviruses, including SL-COVZC45 and RaTG13 [12, 19, 20]. Such viral sequence analysis provides important information regarding genetic characteristics and origin of viruses, and sequence-dependent data can be used for precise diagnosis of etiological agents and adaptation/support of control measures. SARS-CoV-2 dissemination has been reported globally and new infections are recorded with a fast pace in different regions of the world [21]. The growing number of infections over time may result in emergence of new variants. As such, genome sequence tracking and characterization are important to keep track of such events. SARS-CoV-2 sequences phylogenetic analyses will help us to understand the reservoir species, their potential to human transmission and evolution patterns of coronaviruses. The data generated here, where we focus on an in-depth study of SARS-CoV-2 sequences from Saudi Arabia, to further understand the history of this virus.

## METHODS

### Whole genome sequences

GISAID Epiflu Database has a COVID-19 dedicated page (https://www.epicov.org/), from where SARS-CoV-2 genomes are available. We thank the contributors of these sequences (see Acknowledgments, below). The current study intended to compare Saudi SARS-CoV-2 sequences to previously occurring MERS-CoV as well as SARS-CoV and bat-like SARS CoV sequences. Thus, the only three submitted Saudi sequences were used. In addition, a MERS-CoV sequence of Saudi origin, 7 bat SARS-COV sequences collected from 2011 to 2017 and 2 human SARS-CoV sequences were added from NCBI GenBank. Accession number, location and collection dates are shown in Table 1.

**Table 1.** List of genomes used in phylogenetic analysis. hCoV-19 refers to SARS-CoV-2.

| Accession No. | Sample name | Abbreviated name | Data Source | Location | Collection date |
| --- | --- | --- | --- | --- | --- |
| EPI_ISL_416432 | hCoV-19/Saudi Arabia/KAIMRC-Alghoribi/2020 | KAIMRC-Alghoribi | GISAID | Riyadh/Saudi Arabia | 3/7/2020 |
| EPI_ISL_416521 | hCoV-19/Saudi Arabia/SCDC-3321/2020 | SCDC-3321 | GISAID | Riyadh/Saudi Arabia | 3/10/2020 |
| EPI_ISL_416522 | hCoV-19/Saudi Arabia/SCDC-3324/2020 | SCDC-3324 | GISAID | Riyadh/Saudi Arabia | 3/10/2020 |
| MK483839 | MERS_Hu/Albaha-KSA-0800H/2018 | MERS_0800H | NCBI | Albaha/Saudi Arabia | 8/16/2018 |
| MG772933 | bat-SL-CoVZC45 | CoVZC45 | NCBI | Zhoushan city/Zhejiang province/China | 2/2017 |
| KF294457 | bat-SL-CoV_Longquan-140 | Longquan-140 | NCBI | Guizhou province/China | 2012 |
| KY417151 | bat-SL-CoV_Rs7327 | Rs7327 | NCBI | Yunnan Province/China | 10/24/2014 |
| KY417145 | bat-SL-CoV_Rf4092 | Rf4092 | NCBI | Yunnan Province/China | 9/18/2012 |
| KY417142 | bat-SL-CoV_As6526 | As6526 | NCBI | Yunnan Province/China | 5/12/2014 |
| KC881005 | bat-SL-CoV_RsSHC014 | RsSHC014 | NCBI | Yunnan Province/China | 4/17/2011 |
| KP886809 | bat-SL-CoV_YNLF_34C | YNLF_34C | NCBI | China | 5/23/2013 |
| AY278487.3 | Hu-SARS-CoV_BJ02 | BJ02 | NCBI | China | 6/5/2003 |
| AY278489.2 | Hu-SARS-CoV_GD01 | GD01 | NCBI | China | 6/5/2003 |

### Phylogenetic analysis of whole viral genomes

Whole genome alignments were generated by using ClustalW with opening penalty of 15 and extension penalty of 6.66. Pairwise sequence identity and similarity from multiple sequence alignments was calculated using the server (http://imed.med.ucm.es) that contains the SIAS (Sequence Identity And Similarity) tool. Phylogenetic trees were constructed with Neighbor-Joining (NJ) method, Minimum Evolution (ME) method, Maximum Parsimony (MP) method, and UPGMA with 1000 bootstrap replicates (MEGA X) [22].

### Genome recombination analysis

Potential recombination events in the history of the Saudi SARS-CoV-2 sequences were assessed using RDP4 [23]. RDP4 analysis was carried out based on the complete genome (nucleotide) sequence, using RDP, BootScan, GENECONV, Chimera, SISCAN, maximum chi square and 3SEQ methods. These methods are entirely used and compared in order to get consensus results. A putative recombination event was passed to consequent analysis only if it was plausibly defined by at least 3 of the above-mentioned seven algorithms [24]. The minor parent was defined as the one contributing by the smaller fraction of the obtained recombinant, whereas the major parent was that contributing by the larger fraction of the yielded recombinant [25]. Moreover, the recognized recombination events were detected with a Bonferroni corrected *P*-value cut-off of 0.01. In order to avoid the possibility of false-positive results, phylogenetic analysis of the detected recombination was performed [24, 26]. In addition, the whole dataset alignment of each recognized recombinant was divided at the breakpoint positions. If 2 recombination breakpoints existed in a single sequence, the sequence region between the breakpoints was denoted the “minor” region, triggered by the minor parent, while the remaining part is called the “major” region, provoked by the major parent. As a consequence, Neighbor-joining phylogenetic trees were generated to display the probable topological shifts of specific sequences. Phylogenetic discrepancy is revealed by a putative recombinant whose distance in the phylogeny is obviously close to a single parent whilst far from another for each sequence segment [27]. Recombination analysis was repeated for SARS-CoV-2 S gene sequences using automated RDP analysis to investigate the presence of a recombinant that might lead to SARS-CoV-2 emergence among in SARS-CoV-2 sequences.

### Phylogenetic analysis of SARS-CoV-2 S gene sequences

S gene sequences were obtained for 3 Saudi sequences from the GISAID Epiflu Database. In addition, 7 bat SARS-Like CoV sequences, 2 human SARS-CoV sequences and Saudi MERS-CoV sequence were used for alignment. This was followed by finding the best model that could be implemented when constructing the phylogenetic tree upon analysis. Models with the lowest BIC scores (Bayesian Information Criterion) are considered to depict the substitution pattern best. Moreover, AICc value (Akaike Information Criterion, corrected), Maximum Likelihood value (lnL), and the number of parameters (including branch lengths) are considered For each model [28]. Non-uniformity of evolutionary rates among sites may be modeled via applying a discrete Gamma distribution (+G) with 5 rate categories and assuming that a certain fraction of sites is evolutionarily invariable (+I). Furthermore, tree topology was automatically computed to estimate ML values. This analysis involved 13 nucleotide sequences. Evolutionary analyses were conducted in MEGA X [22]. Phylogenetic analysis was performed using the NJ method based on the best fitting substitution model obtained from the previous test with bootstrap of 500 replicates.

### Codon-based Z-test

A codon-based test of positive selection (Z-test, MEGA X) was used to analyze the numbers of non-synonymous and synonymous substitutions per site (dN/dS ratio) in the S gene to check the probability of positive selection occurrences.

### Molecular clock analysis

The molecular clock test was performed by comparing the ML value for the given topology obtained in the presence and absence of the molecular clock constraints under Hasegawa-Kishino-Yano model (+G+I) using MEGA X. Differences in evolutionary rates among sites were modeled using a discrete Gamma (G) distribution and allowed for invariant (I) sites to exist.

## RESULTS

Sequence alignments of whole genomes of SARS-CoV-2, SARS-CoV, bat SARS-like CoVs and MERS-CoV showed an obvious variation in % identity that ranged from extremely high % identities of 99.91-100% identity between Saudi SARS-CoV-2 sequences (suggesting same or similar origin); 78.58-88.03% between Saudi SARS-CoV-2 sequences and bat SARS-like CoVs; 79.18-79.37% between Saudi SARS-CoV-2 sequences and SARS-CoVs that initiated the SARS pandemic in 2003; to relatively low % identity of 52.28-52.3% between Saudi SARS-CoV-2 sequences and Saudi MERS-CoV sequence, as shown in Table 2.

**Table 2.**
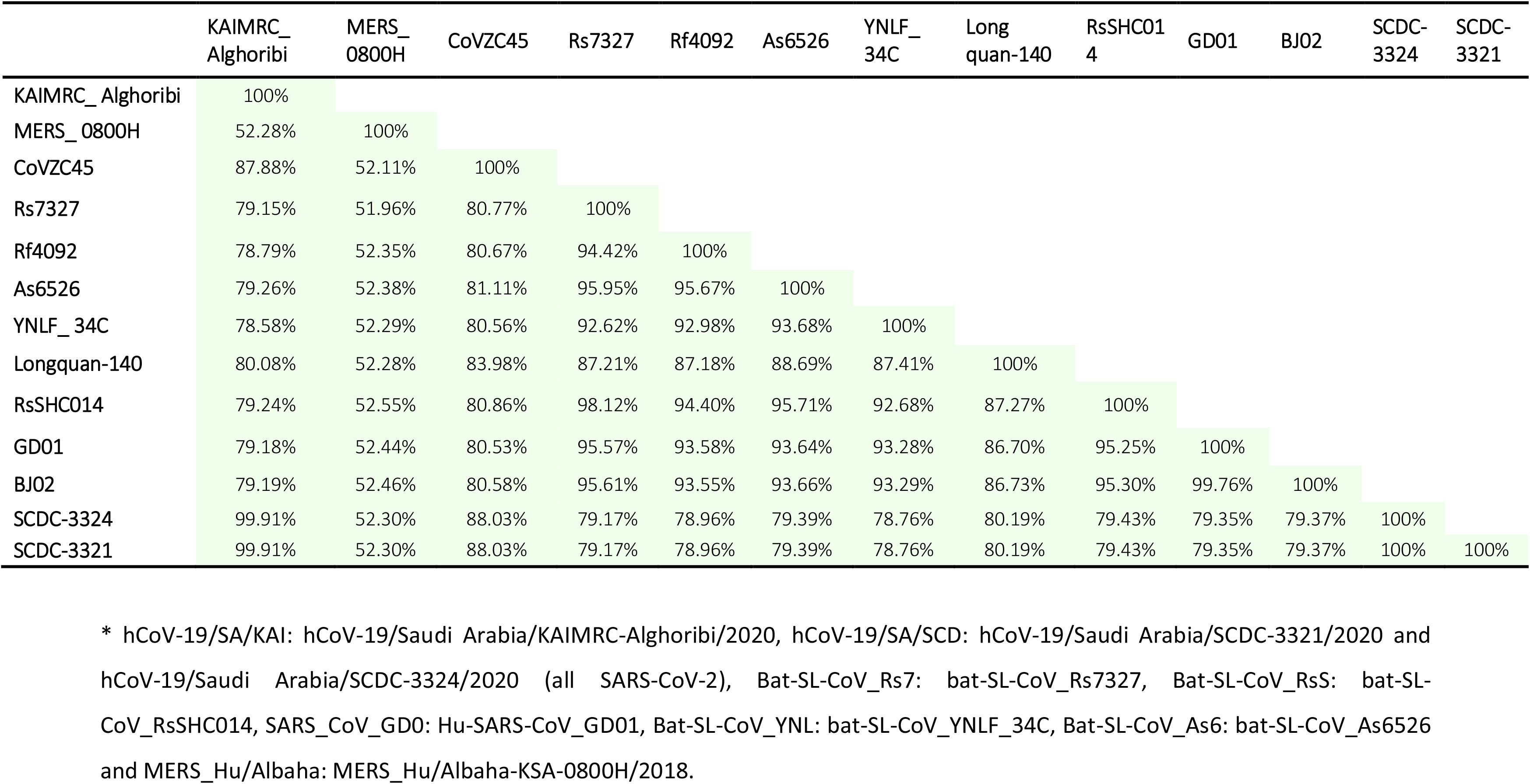
Percent identity between whole genome sequences of studied strains obtained by SIAS (Sequence Identity and Similarity)

Following whole genome alignments, phylogenetic trees were constructed with NJ, ME, UPGMA, and MP methods. The trees had similar topography with significant bootstrap support in case of NJ and ME methods. A tree containing the 3 SARS-CoV-2 Saudi isolates sequences as well as other full-length genomes for the 9 sarbecoviruses of bat and human origin and a merbecovirus, MERS-CoV. Three major clades are observed. The Saudi SARS-CoV-2 isolates form a monophyletic group that nests within a lineage of bat SL-CoVZC45 isolate. This is supported by the percent similarity between the SARS-CoV-2 isolates and bat SL-CoV45 isolate for the full-length genomes (Table 2), which are greater than 87.8%. Eight viruses, 2 human SARS-CoV isolates and 6 bat SARS-like CoV, made up a second distinct lineage and a single MERS-CoV from Abha, a third. In UPGMA, the topology was different, since the monophyletic group comprising the 3 Saudi SARS-CoV-2 isolates was diverged so that it included only 2 isolates, hCoV-19/SA/SCDC-3321 and hCoV-19/SA/SCDC-3324 (100% identity) unlike hCoV-19/SA/KAIMRC-Alghoribi of 99.91% identity to the other 2 SARS-CoV-2. However, the MP method resulted in quite a different phylogenetic topology. Phylogenetic trees generated with each method are shown in Fig 1 and Fig S1. Overall, phylogenetic analysis could reveal that all Saudi viruses with available sequences are of the same or similar origin.

**Fig 1.**
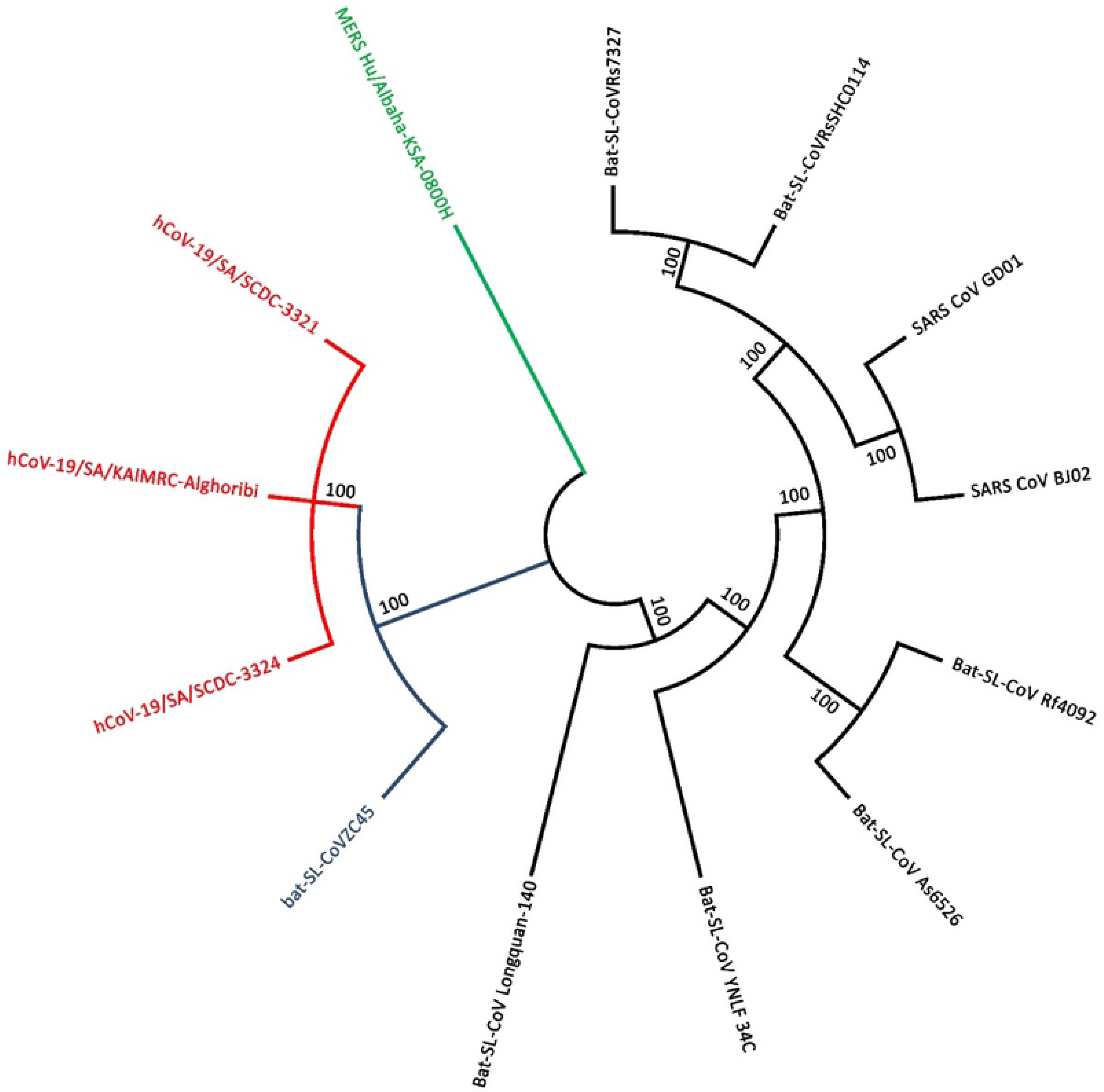
Phylogenetic trees constructed with NJ method to infer evolutionary history using whole genome sequence data of 13 coronaviruses. The bootstrap consensus tree was constructed from 1000 replicates (percentage of replicate trees in which associated strains clustered together are presented at nodes) using MEGA X.

To characterize potential recombination events in the evolutionary history of the sarbecoviruses, the whole-genome sequence of Saudi SARS-CoV-2 and 9 representative coronaviruses— bat-SL-CoVZC45, bat-SL-CoV_Longquan-140, bat-SL-CoV_Rs7327, bat-SL-CoV_Rf4092, bat-SL-CoV_As6526, bat-SL-CoV_RsSHC014, bat-SL-CoV_YNLF_34C, Hu-SARS-CoV_BJ02 and Hu-SARS-CoV_GD01 and MERS-CoV— were analysed using the Recombination Detection Program v.4 (RDP4), in which seven detection methods were used to check each recombinant. MERS-CoV was added to the analysis owing to the coexistence of MERS-CoV in Saudi Arabia (where this virus was first detected) and SARS-CoV-2. Two recombination events were detected between a SARS-CoV-2 (hCoV-19/Saudi Arabia/KAIMRC-Alghoribi) and SARS-like CoVs; these recombination events were also observed for the other Saudi SARS-CoV-2 isolates. The first recombination event was detected by 6 out of 7 detection methods involving RDP, GENECONV, Bootscan, MaxChi, Chimaera & 3SEQ. It included recombination breakpoints at nucleotides 22421 and 22733 which divide the genome into three regions (1-22421, 22422-22732 and 22733-31294) (Fig 2). The major parent of the recombinant was Bat-SL-CoV_YNL34C while the minor parent was Bat-SL-CoV_RsSHC014 as displayed in the recombination event tree (Fig S2). The recombination rate detected was 3.429 × 10^−4^ to 1.102 × 10^−15^ substitutions per site per year at the second region, which comprises the S region. The second recombination event was detected by only 3 detection methods including RDP, Bootscan & 3SEQ. It included recombination breakpoints at 22177 and 22375. The major parent of the recombinant was Bat-SL-CoV_RsSHC014 while the minor parent was Bat-SL-CoV_Rf4 displayed in the recombination event tree (Fig S3). The recombination rate detected was 2.462 × 10^−15^ substitutions per site per year at nucleotides 22134-22217 inside the S region (Fig 3).

**Fig 2.**
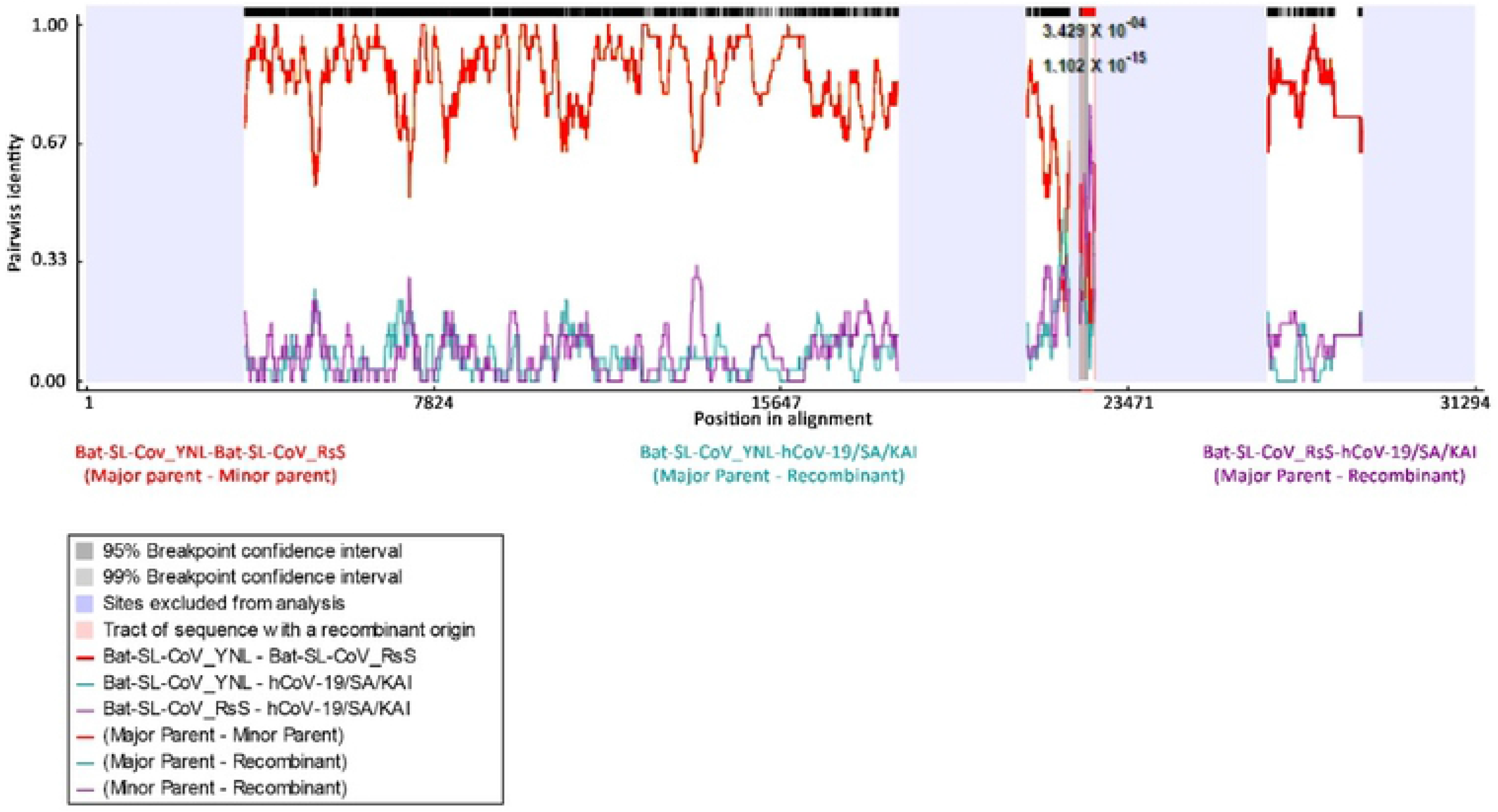
Recombination event 1 in Saudi SARS-CoV-2 isolates. RDP plot reveals two putative recombination breakpoints. The recombination rate is shown at the top. The major and minor parents are shown under the plot. * bat-SL-CoV_RsS: bat-SL-CoV_RsSHC014, bat-SL-CoV_YNLF: bat-SL-CoV_YNLF_34C, hCoV-19/Saudi Arabia/KAI: hCoV-19/Saudi Arabia/KAIMRC-Alghoribi/2020.

**Fig 3.**
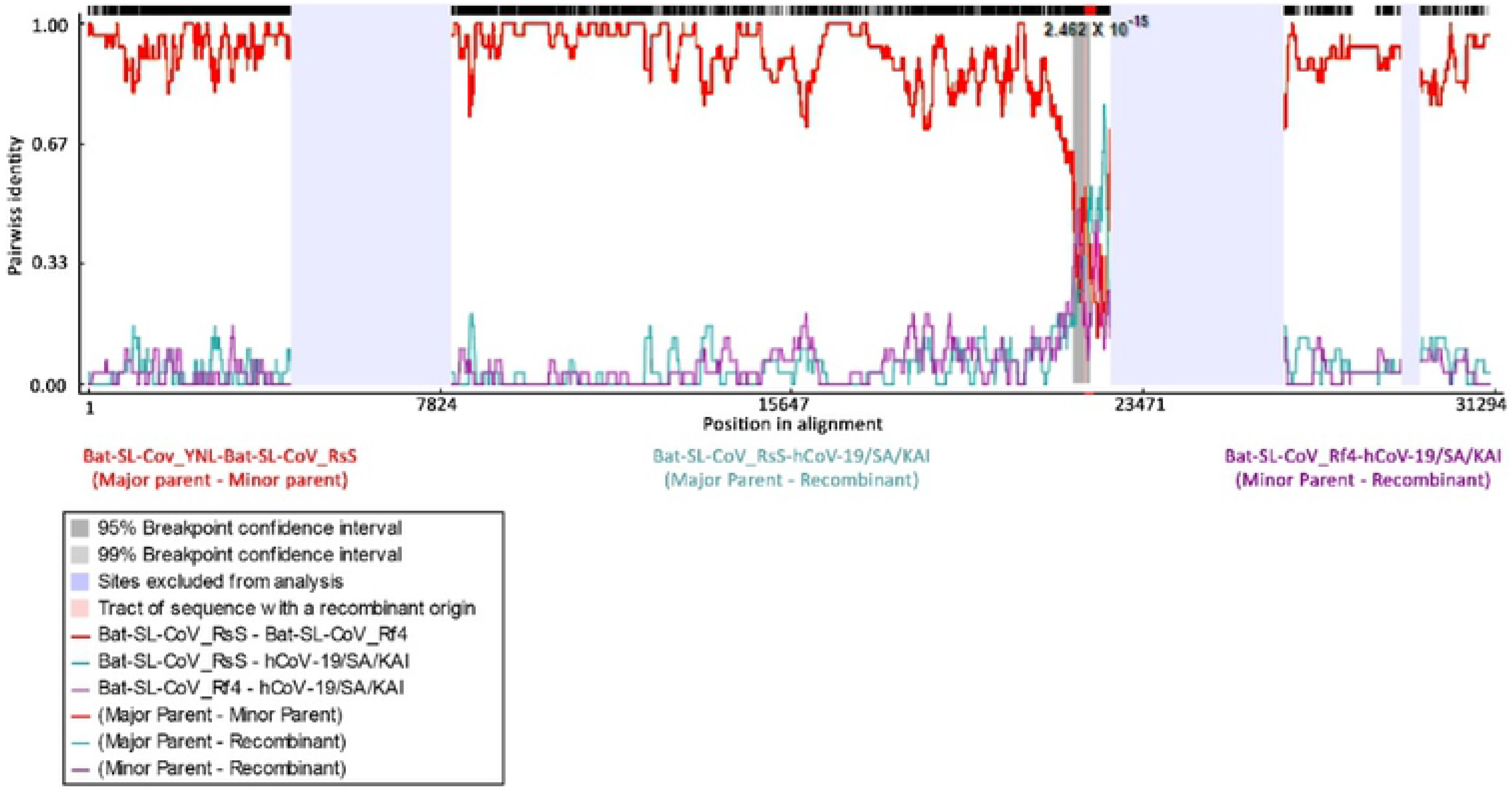
Recombination event 2 in Saudi hCoV-19 isolates. RDP plot reveals two putative recombination breakpoints. The recombination rate is shown at the top. The major and minor parents are shown under the plot. * bat-SL-CoV_RsS: bat-SL-CoV_RsSHC014, bat-SL-CoV_Rf4: bat-SL-CoV_Rf4092, hCoV-19/Saudi Arabia/KAI: hCoV-19/Saudi Arabia/KAIMRC-Alghoribi/2020.

Since both recombination events appeared in the S gene region, sequences of S genes of the 13 CoVs were extracted for multiple alignment using ClustalW, followed by finding the best substitution model to be implemented in the phylogenetic analysis. GTR and TN93 models were the best fitting owing to achieving the least BIC of 51289.85 and 51325.49, respectively. Consequently, the phylogenetic tree was constructed using TN93 model and although, it was constructed using the NJ method (Fig 4), and the obtained tree was consistent with the tree yielded from UPMGA generated previously for the whole genome. Moreover, according to fig 4, bat-SL-CoVZC45 is the closest relative to Saudi SARS-CoV-2 isolates in terms of the S region.

**Fig 4.**
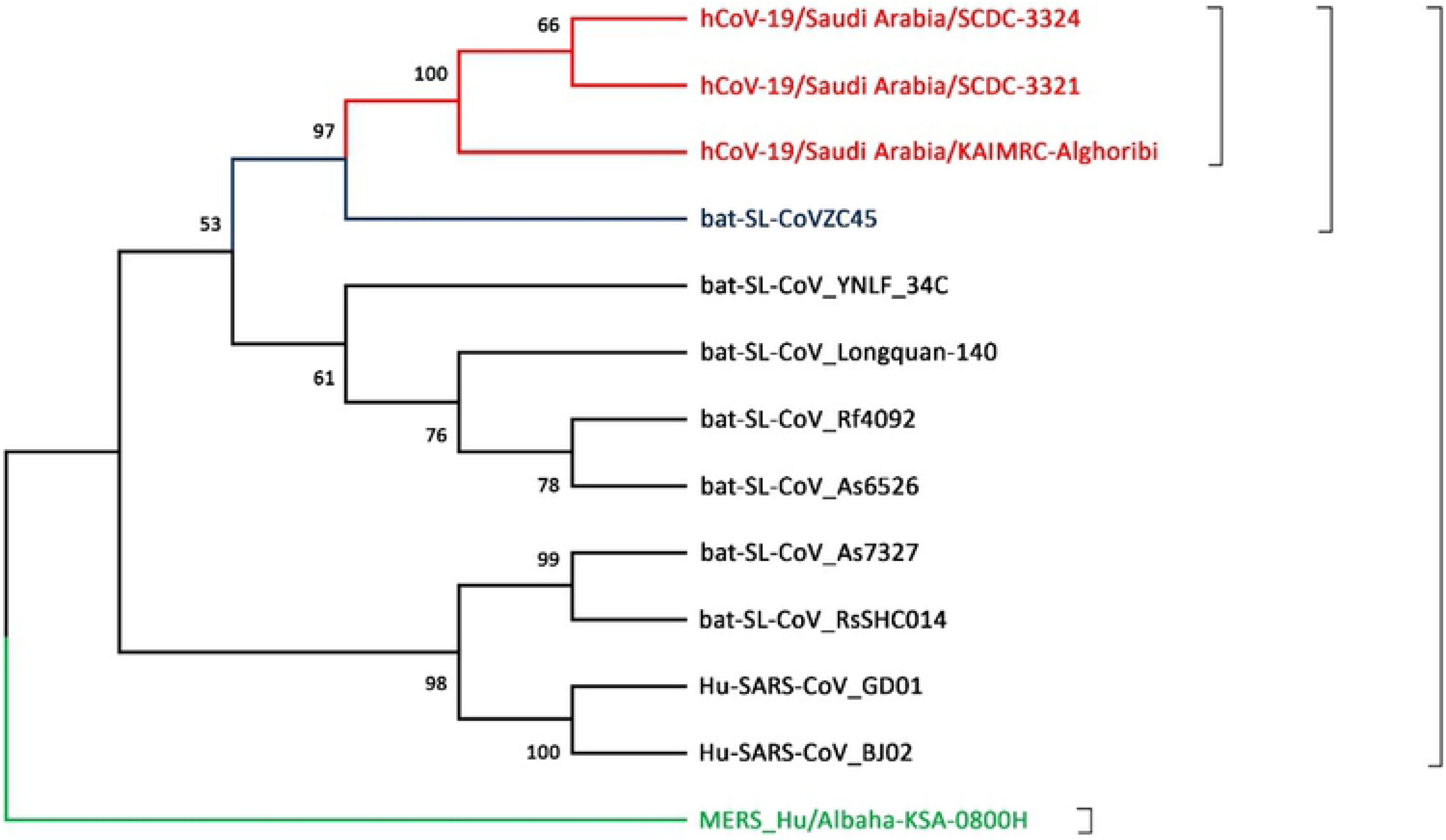
Phylogenetic tree constructed with NJ method using S gene sequence data of the 13 coronaviruses, as described previously. The bootstrap consensus tree was constructed from 500 replicates using MEGA X.

### Positive selection across the SARS-CoV-2 S sequence

To investigate the divergence in sarbecoviruses that may have led to emergence of the novel SARS-CoV-2, positive selection pressure was examined. A codon-based Z-test for positive selection, was used to analyze the numbers of non-synonymous and synonymous substitutions per site (dN/dS ratio) in the S gene. The test showed that positive selection was occurring between the Saudi MERS-CoV_0800H isolate and the bat SARS-like CoV isolates (Bat_SL_CoVZC45, Bat_SL_Rs7327 and Bat-SL_RsSHC014) and the human SARS-CoV isolates (dN/dS =1.7384, 1.9196, 1.7381, 1.89 and 1.8982, respectively, and *P* < 0.0424, *P* < 0.0286, *P* < 0.424, *P* < 0.0306 and *P* < 0.03, respectively; Table 3). However, there was no positive selection observed in the case of the SARS-CoV-2 Saudi isolates (*P* > 0.05). It was proposed that the presence of MERS-CoV strain among the other isolates might have masked any positive selection imposed on SARS-CoV-2 isolates owing to possessing the lowest % identity to the other isolates. Consequently, the codon-based Z-test was carried out again for all isolates except for the MERS-CoV isolate to ensure the proposed hypothesis. It was found that there was positive selection between the Saudi SARS-CoV-2 isolates, bat SL-CoV isolates and human SARS-CoV isolates (*P* < 0.05). The highest positive selection was between Saudi SARS-CoV-2 isolates (hCoV-19/Saudi Arabia/SCDC-3324, hCoV-19/Saudi Arabia/SCDC-3321 and hCoV-19/Saudi Arabia/KAIMRC_Alghoribi) and 2 Bat-SL-CoV isolates (Bat-SL-RsSHC014 and Bat-SL-CoVZC45) (dN/dS = 10.6685, 10.6685, 10.8112, 10.4636, 10.4636 and 10.6251, respectively, and *P* < 0.00001 for all isolates; Table 4), followed by the positive selection between the Saudi SARS-CoV-2 isolates (hCoV-19/Saudi Arabia/SCDC-3324, hCoV-19/Saudi Arabia/SCDC-3321 and hCoV-19/Saudi Arabia/KAIMRC_Alghoribi) and the 2 human SARS-CoV isolates (SARS-CoV_GD01 and SARS-CoV_BJ02) (dN/dS = 8.6491, 8.6491, 8.7746, 8.5216, 8.521 and 8.6457, respectively, and *P* < 0.00001 for all isolates; Table 4). This further suggests that the SARS-CoV-2 isolates are more likely to adaptively evolved from bat SARS-like isolates.

**Table 3.** Codon-based Z-test for positive selection^a^ in the S gene.

|  | MERS_0800H | CoVZC45 | Rs7327 | Rf4092 | As6526 | YNLF_34C | Longquan-140 | RsSHC014 | SCDC-3324 | SCDC-3321 | KAIMRC_Alghoribi | GD01 | BJ02 |
| --- | --- | --- | --- | --- | --- | --- | --- | --- | --- | --- | --- | --- | --- |
| MERS_0800H | - | 1.7384 | 1.9196 | 0.6528 | 1.1736 | 1.2071 | 1.3166 | 1.7381 | 1.6352 | 1.6352 | 1.609 | 1.89 | 1.8982 |
| CoVZC45 | 0.0424 | - | -1.7074 | -3.1417 | -2.0263 | -3.373 | -2.8105 | -1.7738 | -1.0345 | -1.0345 | -1.0668 | -1.7653 | -1.85 |
| Rs7327 | 0.0286 | 1.0000 | - | -2.3669 | -2.9642 | -2.6972 | -3.1513 | -3.4991 | -2.3472 | -2.3472 | -2.3789 | -4.1108 | -4.0906 |
| Rf4092 | 0.2576 | 1.0000 | 1.0000 | - | -5.7237 | -4.1284 | -3.1007 | -3.0026 | -2.9567 | -2.9567 | -2.9890 | -2.9090 | -2.7205 |
| As6526 | 0.1214 | 1.0000 | 1.0000 | 1.0000 | - | -5.2950 | -3.8800 | -3.5282 | -2.3606 | -2.3606 | -2.3925 | -3.4840 | -3.3937 |
| YNLF_34C | 0.1149 | 1.0000 | 1.0000 | 1.0000 | 1.0000 | - | -4.1049 | -3.6478 | -2.3801 | -2.3801 | -2.4120 | -2.5390 | -2.5217 |
| Longquan-140 | 0.0952 | 1.0000 | 1.0000 | 1.0000 | 1.0000 | 1.0000 | - | -4.0380 | -2.1934 | -2.1934 | -2.2253 | -2.7234 | -2.7473 |
| RsSHC014 | 0.0424 | 1.0000 | 1.0000 | 1.0000 | 1.0000 | 1.0000 | 1.0000 | - | -2.7996 | -2.7996 | -2.8313 | -4.2962 | -4.2814 |
| SCDC-3324 | 0.0523 | 1.0000 | 1.0000 | 1.0000 | 1.0000 | 1.0000 | 1.0000 | 1.0000 | - | 0.0000 | 1.0002 | -2.1643 | -2.2197 |
| SCDC-3321 | 0.0523 | 1.0000 | 1.0000 | 1.0000 | 1.0000 | 1.0000 | 1.0000 | 1.0000 | 1.0000 | - | 1.0002 | -2.1643 | -2.2197 |
| KAIMRC_Alghoribi | 0.0551 | 1.0000 | 1.0000 | 1.0000 | 1.0000 | 1.0000 | 1.0000 | 1.0000 | 0.1596 | 0.1596 | - | -2.1960 | -2.2514 |
| GD01 | 0.0306 | 1.0000 | 1.0000 | 1.0000 | 1.0000 | 1.0000 | 1.0000 | 1.0000 | 1.0000 | 1.0000 | 1.0000 | - | 0.4252 |
| BJ02 | 0.0300 | 1.0000 | 1.0000 | 1.0000 | 1.0000 | 1.0000 | 1.0000 | 1.0000 | 1.0000 | 1.0000 | 1.0000 | 0.3357 | - |

<sup>a</sup> Probabilities (*P*) of rejecting the null hypothesis of strict neutrality ( $d_N=d_S$ ) in favor of the alternative hypothesis ( $d_N>d_S$ ) is shown below the diagonal. Values of $P < 0.05$ are considered significant at the 5% level and highlighted. The test statistic values are shown above the diagonal. $d_S$ and $d_N$ are the numbers of synonymous and non-synonymous substitutions per site, respectively. The variance of the difference was computed using the bootstrap method (1000 replicates).
\* hCoV-19/SA/KAI: hCoV-19/Saudi Arabia/KAIMRC-Alghoribi/2020, hCoV-19/SA/SCD: hCoV-19/Saudi Arabia/SCDC-3321/2020 and hCoV-19/Saudi Arabia/SCDC-3324/2020 (all SARS-CoV-2), Bat-SL-CoV\_Rs7: bat-SL-CoV\_Rs7327, Bat-SL-CoV\_RsS: bat-SL-CoV\_RsSHC014, SARS\_CoV\_GD0: Hu-SARS-CoV\_GD01, Bat-SL-CoV\_YNL: bat-SL-CoV\_YNLF\_34C, Bat-SL-CoV\_As6: bat-SL-CoV\_As6526 and MERS\_Hu/Albaha: MERS\_Hu/Albaha-KSA-0800H/2018

**Table 4.** Codon-based Z-test ^a^ of all isolates except for Saudi-MERS-CoV_0800H isolate in the S gene.

|  | CoVZC45 | Rs7327 | Rf4092 | As6526 | YNLF_34C | Longquan-140 | RsSHC014 | SCDC-3324 | SCDC-3321 | KAIMRC_Alghoribi | GD01 | BJ02 |
| --- | --- | --- | --- | --- | --- | --- | --- | --- | --- | --- | --- | --- |
| **CoVZC45** | - | 11.1320 | 7.0741 | 8.3985 | 7.0397 | 8.1521 | 11.3022 | 10.4636 | 10.4636 | 10.6251 | 10.2709 | 10.1542 |
| **Rs7327** | 0.0000 | - | 5.5653 | 6.1583 | 7.4368 | 6.9918 | 3.3205 | 9.7945 | 9.7945 | 9.9286 | 1.0453 | 1.0675 |
| **Rf4092** | 0.0000 | 0.0000 | - | 2.8849 | 5.3215 | 5.6015 | 5.7809 | 6.9788 | 6.9788 | 7.0897 | 6.3999 | 6.7033 |
| **As6526** | 0.0000 | 0.0000 | 0.0023 | - | 5.5098 | 6.8130 | 6.2819 | 8.2467 | 8.2467 | 8.3710 | 5.3357 | 5.4916 |
| **YNLF_34C** | 0.0000 | 0.0000 | 0.0000 | 0.0000 | - | 6.4911 | 7.6653 | 9.6293 | 9.6293 | 9.7592 | 6.8564 | 6.8981 |
| **Longquan-140** | 0.0000 | 0.0000 | 0.0000 | 0.0000 | 0.0000 | - | 7.3144 | 8.2881 | 8.2881 | 8.4126 | 6.7944 | 6.7702 |
| **RsSHC014** | 0.0000 | 0.0008 | 0.0000 | 0.0000 | 0.0000 | 0.0000 | - | 10.6685 | 10.6685 | 10.8112 | 2.6559 | 2.6730 |
| **SCDC-3324** | 0.0000 | 0.0000 | 0.0000 | 0.0000 | 0.0000 | 0.0000 | 0.0000 | - | 0.0000 | -1.0008 | 8.6491 | 8.5216 |
| **SCDC-3321** | 0.0000 | 0.0000 | 0.0000 | 0.0000 | 0.0000 | 0.0000 | 0.0000 | 1.0000 | - | -1.0008 | 8.6491 | 8.5216 |
| **KAIMRC_Alghoribi** | 0.0000 | 0.0000 | 0.0000 | 0.0000 | 0.0000 | 0.0000 | 0.0000 | 1.0000 | 1.0000 | - | 8.7746 | 8.6457 |
| **GD01** | 0.0000 | 0.1490 | 0.0000 | 0.0000 | 0.0000 | 0.0000 | 0.0045 | 0.0000 | 0.0000 | 0.0000 | - | 0.3631 |
| **BJ02** | 0.0000 | 0.1439 | 0.0000 | 0.0000 | 0.0000 | 0.0000 | 0.0043 | 0.0000 | 0.0000 | 0.0000 | 0.3586 | - |

<sup>a</sup> Probabilities (*P*) of rejecting the null hypothesis of strict neutrality ( $d_N=d_S$ ) in favor of the alternative hypothesis ( $d_N>d_S$ ) is shown below the diagonal. Values of $P < 0.05$ are considered significant at the 5% level and highlighted. The test statistic values are shown above the diagonal. $d_S$ and $d_N$ are the numbers of synonymous and non-synonymous substitutions per site, respectively. The variance of the difference was computed using the bootstrap method (1000 replicates).
\* hCoV-19/SA/KAI: hCoV-19/Saudi Arabia/KAIMRC-Alghoribi/2020, hCoV-19/SA/SCD: hCoV-19/Saudi Arabia/SCDC-3321/2020 and hCoV-19/Saudi Arabia/SCDC-3324/2020 (all SARS-CoV-2), Bat-SL-CoV\_Rs7: bat-SL-CoV\_Rs7327, Bat-SL-CoV\_RsS: bat-SL-CoV\_RsSHC014, SARS\_CoV\_GD0: Hu-SARS-CoV\_GD01, Bat-SL-CoV\_YNL: bat-SL-CoV\_YNLF\_34C, Bat-SL-CoV\_As6: bat-SL-CoV\_As6526 and MERS\_Hu/Albaha: MERS\_Hu/Albaha-KSA-0800H/2018.

Next, a molecular clock analysis was carried out using the ML method to examine if the S gene of the 13 isolates used in the current study have the same evolutionary rate throughout the tree. It was found that the strains are not evolving at similar rate indicated by rejection of the null hypothesis of equal evolutionary rate throughout the tree at a 5% significance level (*P* = 0.000E+000) as shown in Table 5.

**Table 5.**
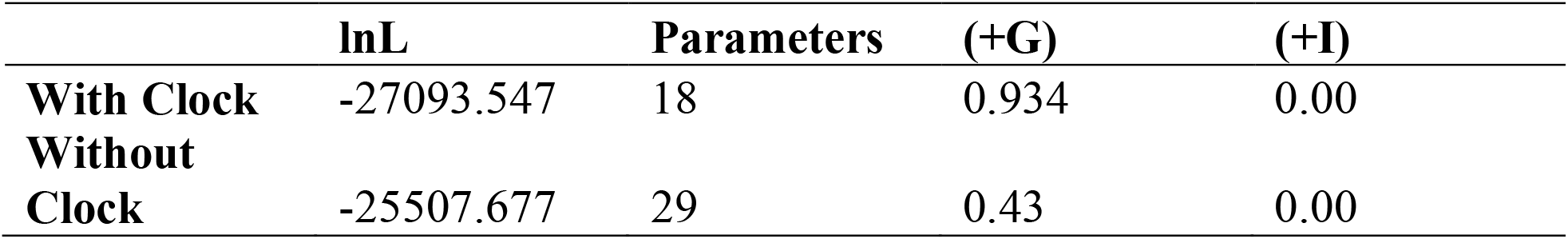
Molecular clock analysis of S gene using the ML method.

The whole genome sequences tested for recombination events using RDP revealed the presence of recombination events in the S region. S gene sequences were checked for recombination events in more details. It was found that two new recombination events have occurred among bat SARS-Like coronavirus, human SARS-CoV (that occurred during the SARS pandemic in 2003) and (SARS-CoV-2) hCoV-19/Saudi Arabia/KAIMRC-Alghoribi S genes; and both recombination events were also observed for the other Saudi SARS-Cov-2 isolates. The first recombination event was detected by 5 out of 7 detection methods involving RDP, GENECONV, Bootscan, MaxChi, SiScan & 3SEQ. It included recombination breakpoints at nucleotides 2094 and 2349 which divide the S sequence into three regions (1-2094, 2095-2348 and 2349-4075). The major parent of the recombinant was Bat-SL-COVZC45 while the minor parent was Hu-SARS-CoV_GD displayed in the recombination event tree (Fig S4). The recombination rate detected by RPD was 1.855 × 10^−3^ substitutions per site per year at 1298-1763 region for all Saudi SARS-CoV isolates (Fig 5), however it increased to 6.039 × 10^−3^ substitutions per site per year when detected by SiScan for hCoV-19/Saudi Arabia/KAIMRC-Alghoribi as a recombinant. The second recombination event was detected by only 2 detection methods including RDP & 3SEQ and was of low quality although the same recombination rate was obtained, however the major parent in the above recombination event was replaced by bat-SL-CoV_As6526.

**Fig 5.**
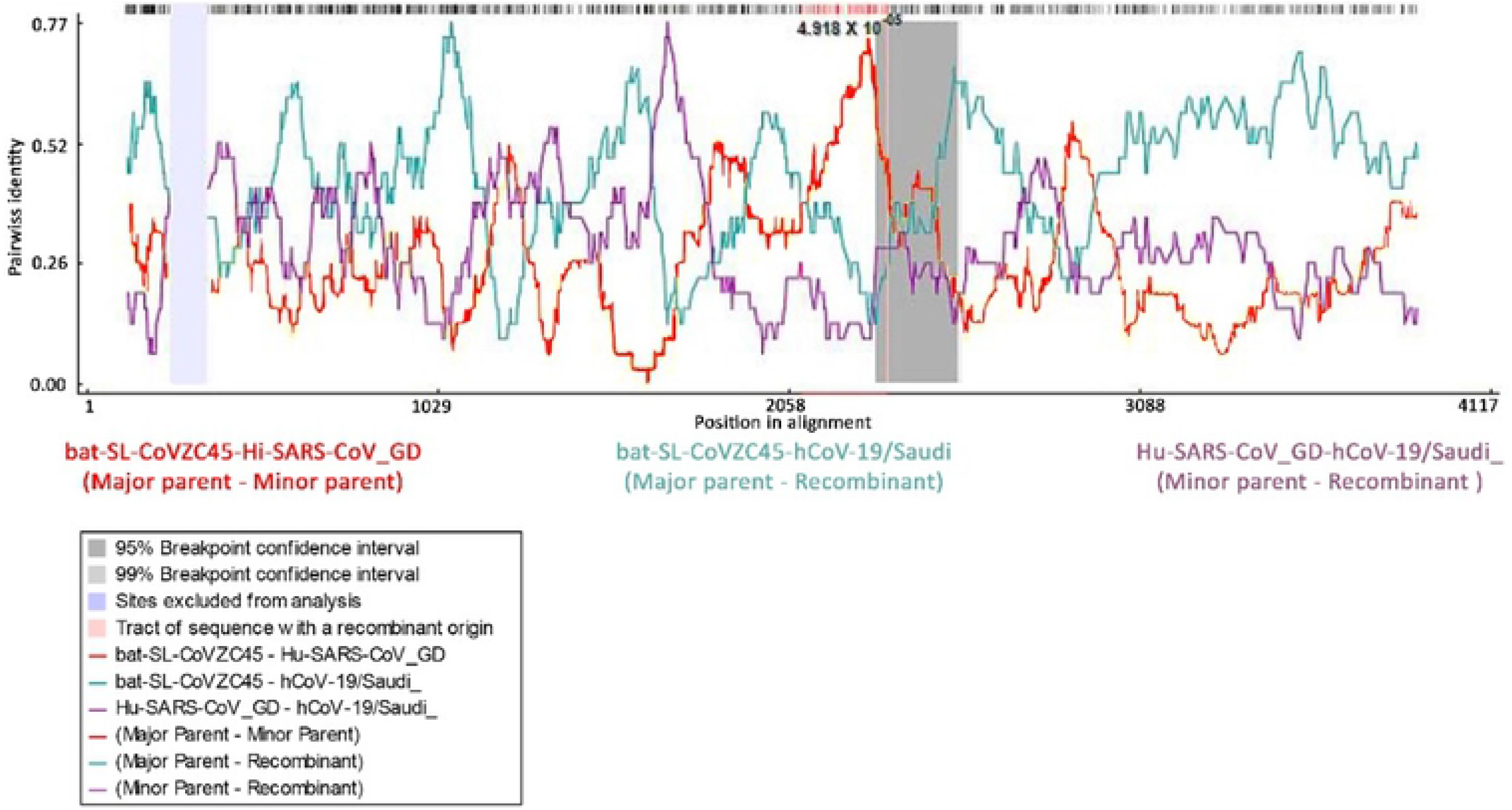
Recombination event in Saudi SARS-CoV-2 S sequences. RDP plot reveals two recombination breakpoints. The recombination rate is shown at the top. The major and minor parents are shown under the plot. * Hu-SARS_CoV_GD: Hu-SARS-CoV_GD01, hCoV-19/Saudi_: hCoV-19/Saudi Arabia/KAIMRC-Alghoribi/2020, hCoV-19/Saudi Arabia/SCDC-3321/2020 and hCoV-19/Saudi Arabia/SCDC-3324/2020 (SARS-CoV-2).

## DISCUSSION

Our knowledge of SARS COV-2 regarding basic and intermediate host species, evolution and genetic variation in relation to other coronaviruses like MERS-COV and SARS-COV is limited. The virus is spreading globally and with an increasing number of infections, its history and evolution needed further investigation. Typically, average evolutionary rate for coronaviruses is roughly considered as 10^−4^ nucleotide substitutions per site per year [29], which agrees with the current study findings.

Phylogenetic analysis of SARS-CoV-2 sequences from Kingdom of Saudi Arabia depicted that they were more similar to bat coronavirus followed by human SARS-CoV, however too distant to MERS-CoV. The results of our phylogenetic analysis are partially in line with a previous study [30, 31], indicative of high similarity with bat SARS-like coronavirus sequences with SARS COV-2 and suggesting that *Rhinolophus* bats may serve as common host for circulating coronavirus. It was previously reported that *Rhinolophus* bats may serve as hosts for potentially emerging viruses [32]. The MP method used for phylogenetic tree designing had a quite different phylogenetic topology from others. This could be owing to the principle of MP method in which the minimum number of evolutionary changes that interprets the whole sequence evolution (tree length) is computed for each topology, and the topology showing the smallest tree length value is chosen as the preferred tree (MP tree) [33]. Although the ME method shares a similar principle, it was mentioned elsewhere that it is closer to NJ method in defining the correct tree and that MP method is less efficient than NJ and ME methods for obtaining the most fitting and/or the correct topology [34]. Consequently, a different topology was expected although it was found in a previous study, that 4 methods led to similar topologies. This may be owing to species differences since the latter was for turkey coronaviruses (group 3 viruses), however the current study was for SARS-CoV-2 virus (group 2 viruses) [35]. Therefore, the suggestion of divergence among Saudi SARS-CoV-2 isolates resulted from MP method was rejected and assumed to be of similar origin.

Recombination events can occur in coronaviruses [36, 37]. As per the present study, recombination analysis of the entire SARS-CoV-2 genome revealed a common isolate in both recombination events which is bat-SL-CoV_RsSHC014, once as a minor parent and another as a major parent. Moreover, the recombination event was detected in the S region. Interestingly, RsSHC014 isolate, which is a bat coronavirus from Chinese horseshoe bats (*Rhinolophidae*) was reported to be significantly more closely related to SARS-CoV than any formerly identified bat coronaviruses, especially in the RBD of the spike protein [38]. This contradicts the findings of a recent study that didn’t recognize any evidence for recombination along the entire genome of SARS-CoV-2 Wuhan isolate [12]. This could be owing to the exclusion of the significant isolate RsSHC014 for whole genome recombination analysis, that can largely limit the identification sensitivity of recombination events of SARS-CoV-2 isolates. Indeed, the exclusion of the significantly putative recombination parent AF531433 influenced the identification sensitivity of recombination events in classical swine fever virus genomes [27].

The current study reported two recombination events between the Saudi SARS-CoV-2 isolates and bat SARS-like CoVs, in the S region, which complements previous suggestions [39, 40]. Such events may relate to the divergence in host tropism. Since the S protein mediates both receptor binding and membrane fusion [40] and is essential for defining host tropism and transmission capacity [41], these sequences were investigated specifically. Recombination events were detected from phylogenetic analysis of S sequences and whole genome. Interestingly, MERS-CoV was found to mask the presence of any positive selection pressure among the Saudi SARS-CoV-2 isolates. This could be due to the distant difference between the two lineages as well as the positive selection pressure sites. Positively selected sites for MERS-CoV are present in the region included the two heptad repeats (HR1 and HR2) and their linker in S2 domain [42], however positively selected sites are located in NTD and RBD of SARS-CoV-2 [43].

Examining the dN/dS ratios for the S gene in Saudi SARS-CoV-2 isolates showed that positive selection was occurring between viruses isolated in 2017 (bat-SL-CoVZC45, Zhoushan city/Zhejiang province/China) and 2020 (hCoV-19/Saudi Arabia/KAIMRC-Alghoribi/2020, hCoV-19/Saudi Arabia/SCDC-3321/2020 and hCoV-19/Saudi Arabia/SCDC-3324/2020, Riyadh/Saudi Arabia) and between viruses isolated in 2011 (bat-SL-CoV_RsSHC014, Yunnan Province/China) and 2020 (hCoV-19/Saudi_Arabia/KAIMRC-Alghoribi/2020, hCoV-19/Saudi_Arabia/SCDC-3321/2020 and hCoV-19/Saudi Arabia/SCDC-3324/2020, Riyadh/Saudi Arabia), after the emergence of the disease in humans. Recombination analysis of S gene and of SARS-CoV-2 whole genome suggested bat SARS-like CoV as the major parental strain. Recombination analysis of S sequence added the possibility of contribution of a SARS-CoV-like sequence though this requires further examination. This finding was supported by a previous study that reported about past recombination detected in the S gene of WHCV (WH-Human 1 coronavirus referred to as ‘2019-nCoV’ of Wuhan, China), SARS-CoV and bat SARS-like CoVs including WIV1 and RsSHC014 isolates [12]. The later isolate agrees with our recombination analysis results obtained for SARS-CoV-2 whole genome. However, recombination analysis of S region revealed another major parental strain which is bat-SL-CoVZC45. This isolate was reported to have a significant nucleotide identity (82.3 to 84%) with SARS-CoV-2 S sequence and is a closer relative [2, 12], and might therefore act as a closer probable ancestor to SARS-CoV-2 [44]. Moreover, the second recombination event, that considered bat-SL-CoV_As6526 isolate as a major parent, was reported to be of low-quality owing to being below the acceptable limit (approved by 2 out of 7 algorithms; minimum approval limit is 3). This might be because of the fact that bat-SL-CoV_As6526 isolate (Betacoronavirus Clade 2) was reported to have deletions in the RBD [45] resulting in enhanced entry using ACE-2 receptor only upon protease treatment, unlike SARS-CoV-2 [46]. However, bat-SL-CoV_As6526 as a recombination contributor is still possible since SARS-CoV-2 S contains most of the contact points with human ACE2 present in clade 1 (Containing SARS-CoV some bat-SL-CoVs as SCH014), besides some amino acid variations which are distinctive to clade 2 (containing the As6526 isolate and other bat-SL-CoVs) and 3 (containing the BM48-31 isolate) [46].

In conclusion, our analysis of 3 Saudi SARS-2-CoV-2 and 7 representative bat SARS-like CoV, 2 human SARS-CoV and MERS-CoV gives further hints about origin of this pandemic virus, in particular with regards to recombination events that underlie SARS-CoV-2 evolution.

## Author contributions

Conceptualization: SE, IN, IOA, AK, AH; data curation: SE; formal analysis: IN, IOA; funding acquisition: SE; investigation: IN, IOA; methodology: IN, IOA; supervision: SE; validation: IN, IOA; writing – original draft: IN, AH; writing – review & editing: IN, IOA, AH, AK, SE.

## Funding information

AK is supported by the UK Medical Research Council (MC_UU_12014/8); SE is supported by KSU Scientific Research Deanship (RGP-VPP-253).

## Acknowledgements

We gratefully acknowledge the authors, originating and submitting laboratories of the sequences from GISAID’s EpiFlu™ Database on which this research is based. The list is detailed below.

---

| Isolate ID | Country | Collection date | Isolate name | Originating Laboratory | Submitting Laboratory | Authors |
| --- | --- | --- | --- | --- | --- | --- |
| EPI_ISL_416521 | Saudi Arabia | 2020-03-10 | hCoV-19/Saudi Arabia/SCDC-3321/2020 | Public Health Laboratory | Public Health Laboratory, Saudi CDC | Albarrag, A |
| EPI_ISL_416522 | Saudi Arabia | 2020-03-10 | hCoV-19/Saudi Arabia/SCDC-3324/2020 | Public Health Laboratory, Saudi CDC | Public Health Laboratory, Saudi CDC | Albarrag, A |
| EPI_ISL_416432 | Saudi Arabia | 2020-03-07 | hCoV-19/Saudi Arabia/KAIMRC-Alghoribi/2020 | Clinical Microbiology Lab | Infectious Disease Research Department, King Abdullah International Medical Research Center (KAIMRC) | Majed Alghoribi, Sadeem Alhayli, Abdulrahman Alswaji, Liliane Okdah, Sameera Al Johani, Michel Doumith |

## SUPPORTING INFORMATION

**Fig S1. Phylogenetic trees constructed with (A) ME, (B) UPGMA and (C) MP methods to infer evolutionary history using whole genome sequence data of 13 coronaviruses.** The bootstrap consensus tree was constructed from 1000 replicates (percentage of replicate trees in which associated strains clustered together are presented at nodes) using MEGA X.

**Fig S2. Phylogenetic tree of recombination event 1 in Saudi SARS-CoV-2 isolates. (A) Phylogenies of the major parental region (1-22421 and 22733-31294) and (B) minor parental region (22422 - 22732). Phylogenies were estimated using UPGMA. The scale bar represents the number of substitutions per site.**

* hCoV-19/SA/KAI: hCoV-19/Saudi Arabia/KAIMRC-Alghoribi/2020, hCoV-19/SA/SCD: hCoV-19/Saudi Arabia/SCDC-3321/2020 and hCoV-19/Saudi Arabia/SCDC-3324/2020 (all SARS-CoV-2), Bat-SL-CoV_Rs7: bat-SL-CoV_Rs7327, Bat-SL-CoV_RsS: bat-SL-CoV_RsSHC014, SARS_CoV_GD0: Hu-SARS-CoV_GD01, Bat-SL-CoV_YNL: bat-SL-CoV_YNLF_34C, Bat-SL-CoV_As6: bat-SL-CoV_As6526 and MERS_Hu/Albaha: MERS_Hu/Albaha-KSA-0800H/2018.

**Fig S3. Phylogenetic tree of recombination event 2 in Saudi hCoV-19 isolates. (A) Phylogenies of the major parental region (1-22177 and 22375-31294) and (B) minor parental region (22178-22374). Phylogenies were estimated using UPGMA. The scale bar represents the number of substitutions per site.**

* hCoV-19/SA/KAI: hCoV-19/Saudi Arabia/KAIMRC-Alghoribi/2020, hCoV-19/SA/SCD: hCoV-19/Saudi Arabia/SCDC-3321/2020 and hCoV-19/Saudi Arabia/SCDC-3324/2020 (SARS-CoV-2), Bat-SL-CoV_Rs7: bat-SL-CoV_Rs7327, Bat-SL-CoV_RsS: bat-SL-CoV_RsSHC014, SARS_CoV_GD0: Hu-SARS-CoV_GD01, Bat-SL-CoV_YNL: bat-SL-CoV_YNLF_34C, Bat-SL-CoV_As6: bat-SL-CoV_As6526 and MERS_Hu/Albaha: MERS_Hu/Albaha-KSA-0800H/2018.

**Fig S4. Phylogenetic tree of recombination event in Saudi SARS-CoV-2 S sequences. (A) Phylogenies of the major parental region (1-2094 and 2349-4075) and (B) minor parental region (2095-2348). Phylogenies were estimated using UPGMA. The scale bar represents the number of substitutions per site.**

* MERS_Hu/Albaha: MERS_Hu/Albaha-KSA-0800H/2018, Bat-SL-CoV_Rs7: bat-SL-CoV_Rs7327, Bat-SL-CoV_RsS: bat-SL-CoV_RsSHC014, Hu-SARS_CoV_GD: Hu-SARS-CoV_GD01, Hu-SARS_CoV_BJ: Hu-SARS-CoV_BJ02, Bat-SL-CoV_As6: bat-SL-CoV_As6526, bat-SL-CoV_Lon: bat-SL-CoV_Longquan-140, Bat-SL-CoV_YNL: bat-SL-CoV_YNLF_34C, hCoV-19/Saudi_: hCoV-19/Saudi Arabia/KAIMRC-Alghoribi/2020, hCoV-19/Saudi Arabia/SCDC-3321/2020 and hCoV-19/Saudi Arabia/SCDC-3324/2020 (SARS-CoV-2).

